# An integrated model of population genetics and community ecology

**DOI:** 10.1101/496125

**Authors:** Isaac Overcast, Brent C. Emerson, Michael J. Hickerson

## Abstract

**Aim:** Quantifying abundance distributions is critical for understanding both how communities assemble, and how community structure varies through time and space, yet estimating abundances requires considerable investment in field work. Community-level population genetic data potentially offer a powerful way to indirectly infer richness, abundance, and the history of accumulation of biodiversity within a community. Here we introduce a joint model linking neutral community assembly and comparative phylogeography to generate both community-level richness, abundance and genetic variation under a neutral model, capturing both equilibrium and non-equilibrium dynamics.

**Location:** Global.

**Methods:** Our model combines a forward-time individual-based community assembly process with a rescaled backward-time neutral coalescent model of multi-taxa population genetics. We explore general dynamics of genetic and abundance-based summary statistics and use approximate Bayesian computation (ABC) to estimate parameters underlying the model of island community assembly. Finally, we demonstrate two applications of the model using community-scale mtDNA sequence data and densely sampled abundances of an arachnid community on La Réunion. First, we use genetic data alone to estimate a summary of the abundance distribution, ground-truthing this against the observed abundances. Then we jointly use the observed genetic data and abundances to estimate the proximity of the community to equilibrium.

**Results:** Simulation experiments of our ABC procedure demonstrate that coupling abundance with genetic data leads to improved accuracy and precision of model parameter estimates compared with using abundance-only data. We further demonstrate reasonable precision and accuracy in estimating a metric underlying the shape of the abundance distribution, temporal progress toward local equilibrium, and several key parameters of the community assembly process. For the insular arachnid assemblage, we find the joint distribution of genetic diversity and abundance approaches equilibrium expectations, and that the Shannon entropy of the observed abundances can be estimated using genetic data alone.

**Main Conclusions:** The framework that we present unifies neutral community assembly and comparative phylogeography to characterize the community-level distribution of both abundance and genetic variation through time, providing a resource that should greatly enhance understanding of both the processes structuring ecological communities and the associated aggregate demographic histories.

## Introduction

The species abundance distribution (SAD) is a classic summary of the structure of ecological communities (McGill *et al*. 2007), which is gaining increasing interest in areas of applied ecology and biodiversity management (Matthews & Whittaker 2015), community assembly (Fattorini *et al*. 2016), and biogeography in general (Matthews *et al*. 2017). However, unbiased comparative species abundance data is often challenging to obtain (Cardoso *et al*. 2011). Standardised sampling protocols can be implemented to improve comparability within studies (e.g. Emerson *et al*. 2017), but these do not account for idiosyncratic phenological or microhabitat differences among species that may affect sampling probability, potentially skewing estimates of relative abundance. Genetic sequence data retains a record of population size changes through time (Griffiths & Tavaré 1994), yet this axis of information has rarely been exploited by community ecologists (Vellend 2005; Laroche *et al*. 2015), and never at the scale of the full community. Therefore, a model linking abundance and effective population size at the community scale could enable a new way to characterize abundance distributions using genetic data alone. Additionally, rather than sampling more individuals to increase resolution of community assembly inference, sampling sequences may allow discrimination between assembly models that are known not to be identifiable with current abundance data alone (Rosindell *et al*. 2012; but see Al Hammal *et al*. 2015). Such rapid and cost effective estimation of SADs could greatly enhance understanding of the structure of ecological communities, with potential to aid in the design of conservation strategies, and to improve forecasts of changes in aggregate population dynamics in the context of global climate change.

The accumulation of sequence data for non-model organisms from over two decades of comparative phylogeographic studies (Avise *et al*. 2016), large-scale DNA barcoding initiatives (e.g. Bucklin *et al*. 2011), and forthcoming community-scale genome-wide data (Garrick *et al*. 2015), presents an exciting opportunity for linking abundances and aggregate population genetic data. However, we lack a flexible joint model that links existing models in comparative phylogeography (Satler & Carstens 2017; Xue & Hickerson 2017) with existing biogeographic models of community assembly (*Rosindell et al*. 2012; Rosindell & Harmon 2013).

Despite the potential of comparative phylogeography to leverage the power of aggregated demographic histories to answer fundamental questions about community assembly and macroecology (Avise et al. 1987; Hickerson *et al*. 2010; Avise *et al*. 2016), such approaches have generally neglected the growing body of theory from community ecology that seeks to accommodate the relative importance of deterministic (Tilman 2004; Maire *et al*. 2012) and stochastic processes (MacArthur & Wilson 1963; Hubbell 2001; Rosindell *et al*. 2012) governing the assembly of communities. For instance, comparative phylogeographic approaches that do incorporate community assembly have tended to focus on general models of shared demographic histories (Burbrink *et al*. 2016; Satler & Carstens 2017), rather than models that are explicitly parameterized from ecological community assembly theory (but see Bunnefeld *et al*. 2018).

Ecological theory has been fundamental for understanding processes underlying spatial patterns of biodiversity as typically quantified by regional SADs and species area relationships (McGill *et al*. 2007). However, ecological models of community assembly tend to view communities as static pools with an ahistorical focus on equilibrium expectations (Weiher *et al*. 2011). Although there have been efforts to incorporate non-equilibrium history in models of community assembly (Clark & McLachlan 2003), as well as a long tradition of incorporating phylogenetic information (Webb *et al*. 2002; Jabot & Chave 2009) at also accommodates non-equilibrium historical dynamics (Pigot & Etienne 2015; Manceau *et al*. 2015), there has only been limited, yet promising, effort in considering intraspecific genetic polymorphism within a dynamic non-equilibrium assembly framework (Vellend *et al*. 2014; Laroche *et al*. 2015) or within statistical models of macro-ecology (Smith *et al*. 2017; Pelletier & Carstens 2018). These efforts in bridging the gap between ecological models and population genetics have focused on characterizing the correlation between species diversity and genetic diversity in ecological communities (the species-genetic diversity correlation: Vellend 2005; Papadopoulou *et al*. 2011; Vellend et al. 2014, Laroche *et al*. 2015) while other efforts have looked at the relationships between adaptive genetic diversity and community dynamics (Hughes *et al*. 2008; Becks *et al*. 2010; Schoener 2011).

Despite these important efforts to unify our understanding of ecological and evolutionary dynamics, a community-scale model linking species abundances and genetic diversities under a dynamic model of assembly has yet to be proposed. Here we describe, test, and demonstrate a joint inferential framework that bridges ecological neutral theory with population genetics in order to make joint predictions of community-wide distributions of species abundances, genetic diversities, and genetic divergences under a dynamic non-equilibrium model of assembly. The unified framework we present combines a forward-time model of community assembly with a backward-time coalescent model, linking abundance and colonization history with aggregated population genetic samples from multiple taxa.

We use simulation experiments to validate the power and accuracy of our method using an approximate Bayesian computation framework (ABC; Csilléry *et al*. 2012) for estimating model parameters. Similar to Jabot & Chave (2009), who used phylogenetic information to estimate parameters of a neutral community assembly model with ABC, we merge population genetics and a similar neutral ecological model in an ABC context. After using simulations to validate the method, we demonstrate an application to a sample of community-wide mitochondrial DNA sequence data and corresponding densely sampled abundance estimates obtained from an assemblage of 57 spider species from the island of Réunion (Emerson *et al*. 2017). Using only the sequence data, we accurately estimate the Shannon entropy of the observed SAD, and additionally obtain an estimate of the equilibrium state of the community. The joint model, implemented in Python, and all ipython notebooks for reproducing simulations and analysis are freely available on GitHub: https://github.com/isaacovercast/gimmeSAD.

## Materials and Methods

### Model Overview

First, forward-time community assembly simulations are performed using an island/mainland metacommunity model following Rosindell & Harmon (2013). This individual-based neutral model unifies MacArthur and Wilson’s equilibrium theory of island biogeography (ETIB) with Hubbell’s unified neutral theory of biodiversity (UNTB) to generate time-dependent non-equilibrium and equilibrium predictions of local richness and abundances (MacArthur & Wilson 1963; Hubbell 2001). We use these predicted temporal changes in abundance distributions and colonization times to parameterize a multi-species model of aggregate population genetic data backwards in time under the coalescent (Rosenberg & Nordborg 2002). The former allows for inference about the time series progression of community change while the latter links predicted changes in community population genetic data to this community assembly process (see Fig. 1 & Box S1).

**Figure 1.**
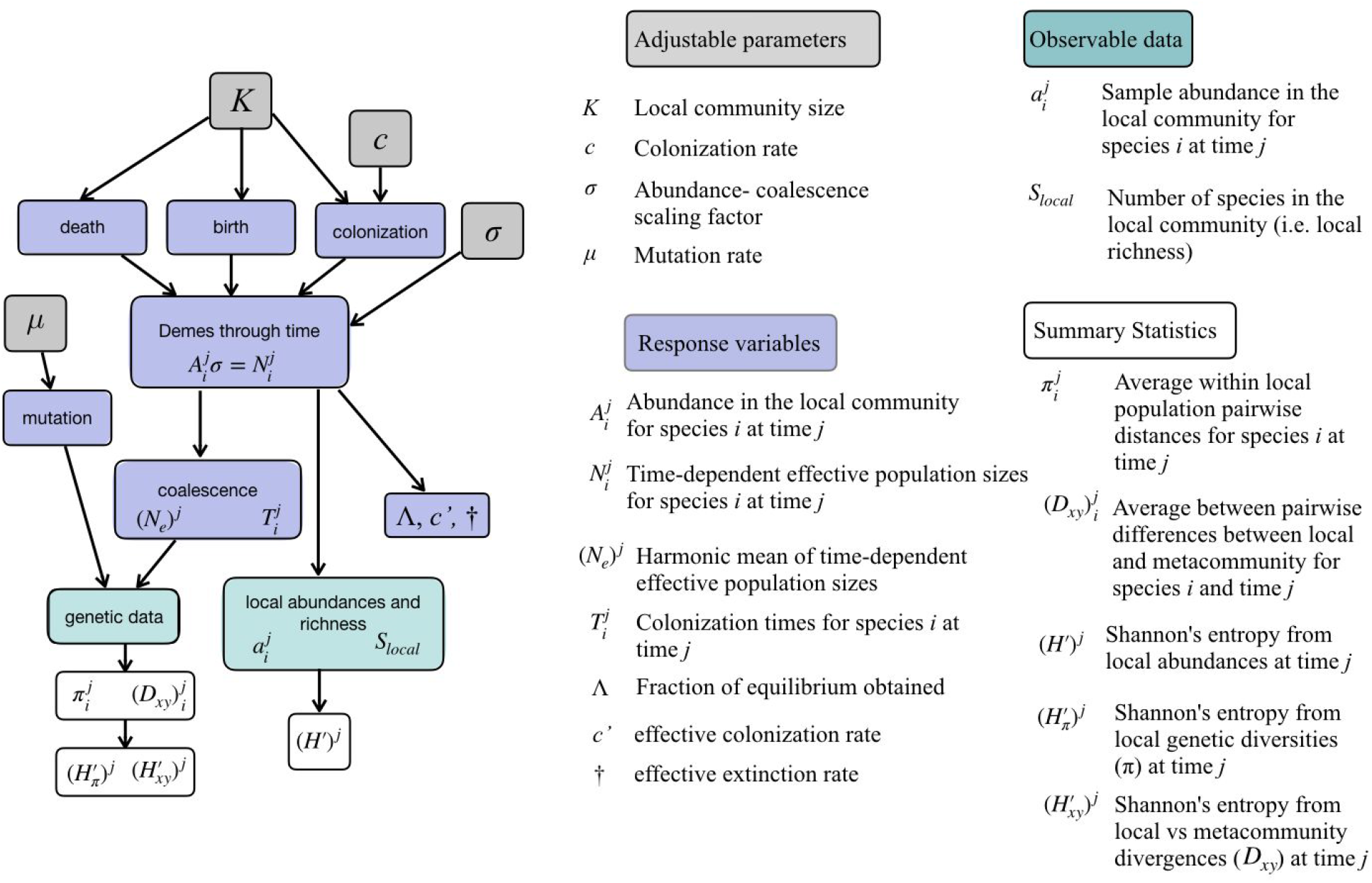
Schematic version of the model parameters, processes, and response variables. Directed acyclic graph (DAG) depicting the conditional dependencies underlying the joint model’s set of adjustable parameters, response variables, data, and associated summary statistics (under model configuration *M_AMI_*). For simplicity, the components of the source metacommunity are elided from the figure, but are described fully in the text.

### Forward-Time Model

Forward-time simulations of community assembly follow the spatially implicit neutral model of Rosindell & Harmon (2013) that unifies the ETIB with the UNTB whereby abundance distributions, and immigration and extinction rates proceed under a birth/death/colonization process in the biogeographical context of a focal local community and a regional source pool (metacommunity). In this model the carrying capacity (K) of the local community is fixed and of finite size. The colonization rate is modeled as a single parameter (*c*) that specifies the probability of a colonization event. Colonizing species are sampled from a metacommunity composed of species with abundances that are independently and identically distributed according to the logseries distribution (Fisher *et al*. 1943), and which is static with respect to the timescale of local assembly. At each time-step one individual is randomly sampled for removal from the local community. With probability 1 - *c*, this individual is replaced by the offspring of a randomly sampled individual from the local community. With probability c, the individual is replaced by a randomly sampled member of the metacommunity, where the probability of sampling from any given species is weighted by the relative metacommunity abundance (*A_meta_*; Table S1).

Given that the forward-time process follows a Moran model, we will refer to one birth/death event as a time-step, with one generation encompassing *K* time-steps. Information about the state of the community in the forward-time simulation model is recorded at regular time-intervals of arbitrary length, with the default interval length being equal to 100000 time-steps. Model state can be described by a vector of *T_i_* = {τ_1_,…, τ*_S_local__*} containing the time since colonization (in generations) for species *i* in the local community as well as a jointly associated vector *A_i_* = {*A*_1_,…, *A_S_local__*} that contains the associated abundances for species *i* in the local community. The history of abundance changes for species *i* going back in time τ_*i*_ generations is contained by a vector 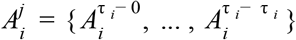 such that *j* = τ_*i*_ – 0 decreases going back in time at prior time intervals until τ_*i*_ – τ_*i*_. The counts of post-colonization migration events are accumulated per species in the vector *M* = {*m*_1_,… *m_S_local__* } (Table S1). Two emergent pseudo-parameters (model response variables) are then c′ (effective colonization rate) and † (effective extinction rate) which are defined as the realized number of colonization and extinction events per generation, respectively (Table S2).

### Scaling forward-time model to backward-time coalescent model

For the *i*-th local species that is extant at the *j*-th time interval with an abundance of 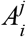, there exists the history of changes in abundance over time since colonization *τ_i_*, from a source species in the metacommunity. To relate raw sample-based abundances with the effective population sizes that parameterize the backward-time coalescent process of the gene tree lineages, we make the assumption of a random spatial distribution of individuals that is predicted to lead to a simple scaling relationship whereby the sample-based and regional-based abundance distributions have the same functional form (Green & Plotkin 2007). To approximate this expectation, we incorporate a rescaling that is based on the assumption that the observed local abundances from direct sampling are proportional to regional abundances and thus current effective population sizes.

To this end we rescale the time-dependent abundance of each species 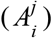 into a time-dependent effective population size 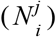 using the scaling factor o such that 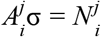. This is equivalent to assuming the abundance counts 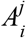 actually record the number of demes, each of size σ, over time per species. Across all species sampled genetically at the *j*-th time interval, this yields time dependent vectors of the effective population sizes for species 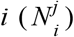, the associated times since colonization in units of generations *T_i_* = {*τ*_1_,…, *τ_S_local__*}, and temporally static effective population size vectors for the corresponding source metacommunity species (*N_meta_*). Under this assumption, each local species consists of a metapopulation consisting of 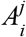 demes of size σ with strong migration conditions that reduce to the temporally dynamic predictions of a panmictic effective population of size 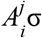. Under this assumption, the “collecting phase” is predicted to dominate the entire history of ancestry thereby approaching the standard panmictic coalescent expectations of a time dependent effective population size 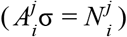 as the number of demes become large (Wakeley & Aliacar 2001; Wakeley 2001; Wakeley 2004). Importantly, this rescaling is based on the assumption that the observed abundances from direct sampling are proportional to actual abundances and current effective population sizes, although these relationships are known to be complex (Luikart *et al*. 2010). However, how σ changes the timescale of both forward and backward processes is not determined in our model and therefore it is critical to determine if a chosen σ value results in a model that can generate the observed data. As a check, one should assess the ability of the model to generate the data by statistical goodness of fit tests or model evaluation (Gelman 2003; Lemaire *et al*. 2016). Alternatively, o could be treated as an unknown parameter and estimated given the data.

Given the parameters of the backward-time model (Tables S1 & S2), we use the *msPr7me* coalescent simulator program (Kelleher *et al*. 2016) to generate genetic polymorphism data matching an arbitrary sampling regime of the local and/or metacommunity species pair sample sizes (with respect to numbers of individuals sampled at a mtDNA locus of length L). Instead of parameterizing the coalescent simulations of the 7th species following the 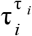 stochastic changes in effective population sizes since colonization according to 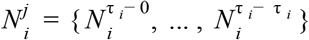, we use *N_e_i__*, the harmonic mean of each species’ effective population size across all time steps indicated by the 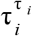 elements within 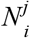 (Karlin 1968; Pollak 1983). One gene genealogy is simulated for each sampled species pair corresponding to a 570bp segment of the mitochondrial COI gene, and mutations are applied under an infinite-sites model given an assumed invertebrate mitochondrial divergence rate (1.1% per species per million years; e.g. Brower 1994).

### Initial Conditions

We implement two different starting conditions to simulate volcanic versus continental island formation. Our initial conditions under the volcanic model deviate from those of Rosindell & Harmon (2013), in that at time zero they assume that one initial colonizing lineage consumes all available space in the community, thereby saturating *K*. In our model we select the most abundant species in the metacommunity and introduce one individual into the unpopulated local community. This initial condition is both biologically more realistic, and also avoids the assumption that volcanic island carrying capacity is saturated at time zero, which could generate unrealistic quantities of genetic diversity in the initial colonizing lineage. Continental islands are initially populated by making *K* independent random samples from the metacommunity proportional to their relative abundances. Here we are modelling a community of panmictic species that are simultaneously and instantaneously isolated from the metacommunity at time zero. Because we assume panmixia prior to isolation, the vector of colonization times 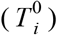 are initially identical across the entire local community. Following subsequent local extinction and replacement by new colonizing species, the vector of colonization times 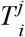, becomes heterogeneous.

### Quantifying Equilibrium

Equilibrium is commonly defined as the dynamic balance between colonization and extinction rates that emerges over time, eventually leading to a stationary distribution where the two rates are expected to be equal (MacArthur & Wilson 1963). However, under certain conditions, species richness and abundances may fail to equilibrate simultaneously, in which case the classic definition of equilibrium is insufficient (see Rosindell & Harmon 2013). To address the need for a more robust concept we follow Rosindell & Harmon (2013) in defining equilibrium as the point at which the starting conditions of the model are no longer detectable in the state of the system. In addition to colonization/extinction rate balance, this auxiliary definition guarantees that both richness and the SAD have reached their expected equilibrium values. Here we define a new term to measure the fraction of this equilibrium obtained by the community and treat it as an emergent parameter that can be estimated by sampling the prior and posterior distribution (Λ; Table S2). This quantity is defined as 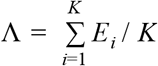, where *K* is the carrying capacity and *E* is the boolean vector of length *K* such that *E_i_* for *i* = {1,…, *K*} indicates the colonization status of each individual in the local community. The value of Λ therefore ranges from 0 to 1 with small values indicating early assembly history, and larger values indicating later assembly history and approach to equilibrium. When all individuals present in the local community are descended from a lineage that colonized during the simulation then 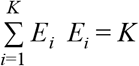 and Λ = 1. Our model of community assembly is inherently stochastic, so the amount of time for any given simulation to reach equilibrium is a random variable given the distribution under the model. For each forward-time simulation we track elapsed time, local community composition (both abundances and richness), and colonization times for all local species. We are interested in equilibrium and non-equilibrium dynamics, so we poll this information at regular intervals of arbitrary duration.

### Summary Statistics

At each time interval we extract the simulated sequences from a sample of the local community and calculate nucleotide diversity as the average number of pairwise differences (π; Tajima 1983) within the local community for each sampled species given *S_local_* (*π_i_* = {*π*_1_,… *π_n_*}). We then summarize the distribution of genetic variation by constructing a one dimensional histogram (Y) of local community genetic diversity such that:

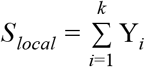

where *k* is the number of bins (with *k*=10 for all simulation and empirical analyses), and bin width max(*π_i_*)/*k*. We term this summary of local community diversity the one dimensional species genetic diversity distribution (1D-SGD; Fig. S1). Next, we calculate absolute divergence (D_xy_; Nei 1987) between samples from each local-metacommunity sister pair (D_xy_i_ = {D_xy_1_,… D_xy_n_}). The values of *π_i_* and D_xy_i_ are aggregated across all species-pairs sampled from the community within each time-point and summarized as a *k* × *k*joint frequency histogram (*X*) with equal-width bins such that:

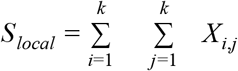

The upper bound for each dimension of the histogram is fixed to the maximum values of *π* and D_xy_ within a given simulation. We term this joint summary of community diversity/divergence as the two dimensional species genetic diversity distribution (2D-SGD; Fig. S2). The 1D and 2D-SGD are simple histograms that collapse the full distribution of community genetic diversity into a summary representation. Additionally, at each time interval we record the rank abundance curve (RAC), the SAD, and Shannon entropy calculated for the community (H’; Boltzmann 1872; Shannon 1948; Hill 1973), which is:

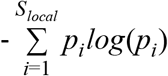

where *p_i_* is the proportional abundance of species *i*. Given an observed sample of *S_local_* species sampled from an empirical community, the simulated summary statistics are filtered to match the observed sampling configuration. As an additional method of comparison with the H’ derived from the SAD, we also calculated the Shannon entropy derived for both the 1D-SGD (*π*), and distribution of D_xy_ per sampling time point and notate this as H’_π_ and H’_Dxy_ respectively.

### Simulation Study Design

To characterize the joint temporal dynamics of the SAD and 2D-SGD under non-equilibrium and equilibrium community assembly, we simulated assembly histories for both continental and volcanic islands, under a range of parameter values using a range of local community sizes (*K* = 1000, 5000, 10000) and colonization rates (*c* = 0.0001, 0.001, 0.01). We generated 10,000 replicated simulations for each combination of origin type, local community size, and colonization rate, resulting in 180,000 total simulated community histories. All forward-time simulations were run for twice the mean time to turnover equilibrium (Λ) for the largest local community with the smallest colonization rate (5 × 10^9^ generations). We then summarized the temporal changes in H’, π, D_xy_, H’_π_, and H’_Dxy_ by calculating the mean and standard deviation of each of these metrics for each parameterization across replicate sets of simulations at five values of Λ (0.1, 0.25, 0.5, 0.75, 1). For this initial set of exploratory simulation experiments, we calculated H’ on the entire set of species while π, D_xy_, H’_π_, and H’_Dxy_ were likewise calculated on this entire set of *S_local_* species given samples of 10 individuals per species in the local community and associated metacommunity source populations.

### Bias and Accuracy in Estimating Parameters

Next, we evaluated the suitability of H’ and the relative bin magnitudes of the SGD as summary statistics for parameter estimation using ABC by conducting a suite of leave-one-out cross-validation experiments under various configurations (Table 1) whereby parameters of known values are estimated (Csilléry *et al*. 2012). We focus on evaluating accuracy and precision in estimating the following community-wide model parameters and pseudo-parameters: local community size (*K*), parameterized colonization rate (*c*), fraction of equilibrium (Λ), realized colonization rate (*c′*), extinction rate (†), and Shannon entropy (H’). We additionally explored estimation of community-wide parameters given various sequence and abundance data availability configurations (see Table 1). For example, given only the DNA sequence data sampled from a focal local community, the relative bin magnitudes of the observed 1D-SGD can be used as the summary statistic vector and both H’ and Λ can be estimated, along with the other model parameters such as *c*, and † (ABC configuration *M_I_*; Table 1).

**Table 1.** ABC Model Configurations. An overview of the five different model configurations explored, indicating summary statistics derived from available observed data (π, D_xy_, and/or H’) and the parameters to be estimated under our ABC framework. These scenarios arise from various combinations of observed abundances (A), community-scale nucleotide diversity (I), and local-metacommunity divergence (M). The Shannon entropy (H’) can be configured either as a summary statistic or as an estimated pseudo-parameter, depending on whether densely sampled abundances are available for the community of interest. Other pseudo-parameters (*c, c′, K*, †, Λ) can be estimated under all ABC configurations.

| Model Configuration | Summary Statistic Vectors |  |  | Estimated Pseudo-Parameters |
| --- | --- | --- | --- | --- |
| $M_A$ | -- | -- | $H'$ | $c$, $c'$, $K$, †, $\Lambda$ |
| $M_I$ | $\pi$ | -- | -- | $c$, $c'$, $K$, †, $H'$, $\Lambda$ |
| $M_{AI}$ | $\pi$ | -- | $H'$ | $c$, $c'$, $K$, †, $\Lambda$ |
| $M_{MI}$ | $\pi$ | $D_{xy}$ | -- | $c$, $c'$, $K$, †, $H'$, $\Lambda$ |
| $M_{AMI}$ | $\pi$ | $D_{xy}$ | $H'$ | $c$, $c'$, $K$, †, $\Lambda$ |

To construct the reference table for the cross-validation analyses, we performed 1,000,000 community assembly simulations, sampling parameter values of *K, c*, and Λ according to uniform prior distributions (*K* = ~U(1,000-10,000), *c* = ~U(0.0001-0.01), and Λ = ~U[0, 1); see Table S1 for all simulation parameters). We then conducted ABC leave-one-out cross-validation using the *cv4abc* function of the *abc* R package (Csilléry *et al*. 2012). For the ABC procedure we used simple rejection sampling and a tolerance sufficient to retain 1000 samples from the prior to construct the posterior estimate for each parameter of interest. We performed 100 leave-one-out cross-validation replicates per data configuration for each estimated parameter, and quantified accuracy of parameter estimation by calculating root-mean-square error (RMSE) and the coefficient of determination (R^2^) for sampled and estimated parameter values.

### Empirical Application

Following our simulation experiments demonstrating that the ABC model can effectively estimate parameters, we perform an empirical analysis on a published dataset from a community of spiders from the island of Réunion, an overseas department of France located in the Indian Ocean approximately 900 km east of Madagascar. In the original study, using a standardized protocol spiders were sampled from 10 lowland rainforest plots distributed across the island and sorted into 57 presumed biological species using a protocol combining morphological sorting and mtDNA sequencing (570bp Cytochrome Oxidase c Subunit I; Emerson *et al*. 2017). The dense and standardized sampling allows us to use both the H’ calculated from the observed SAD as well as the 1D-SGD calculated from the observed sequence data for estimating assembly model parameters. Therefore, we use model configuration *M_I_* to estimate H’, and *M_A_, M_I_*, and *M_AI_* to alternatively estimate Λ (Table S1). Under all model configurations we estimate parameters c’ and †. For the ABC inference procedure, we simulated 1,000,000 samples by drawing parameter values from the same prior distribution used for the cross-validation analysis, and used the same rejection method to accept the closest 1,000 data sets to sample from the posterior distribution. When calculating n for each island taxon we used sample sizes with respect to numbers of individuals matching the observed spider data exactly with respect to numbers of individuals and length of DNA sequence. Simulations for the empirical analysis were run on a 40 core Intel Xeon 2.20GHz workstation with 256GB of main memory and were completed in approximately 1 week.

We evaluated the overall goodness of fit of our posterior estimate to the observed data in two ways. First, we quantified the absolute Euclidean distances between the retained and observed summary statistics. Additionally, we performed a prior predictive check by projecting the retained simulated SGD, along with the observed SGD into principal component (PC) space. A good fit of the model to the data should generate simulated summary statistics sufficiently similar to those of the observed data as to be indistinguishable in the PC analysis.

## Results

### The Joint SAD and SGD Through Time

The classically lognormal-like shape of the SAD, with most species of low abundance, is mirrored by the distribution of genetic diversities (Fig. 2). The shape of the joint spectrum of community genetic diversity (π) and genetic divergence (D_xy_) generally widens over time as richness increases, while the corresponding H’ of the SAD generally increases over the same time intervals (Figs. 2 & 3). We find that most species display low amounts of standing genetic diversity, as characterized by average pairwise differences (π), although there are important temporal dependencies as these characteristics only accrue with time as A progresses. On the other hand, time has a reduced impact on the distribution of D_xy_, which obtains the lognormal-like shape even at very early stages of assembly, although with greater variability, as expected given that the final waiting times in the larger ancestral population will predict a large variance in this summary statistic, regardless of colonization time (Takahata & Nei 1985).

**Figure 2.**
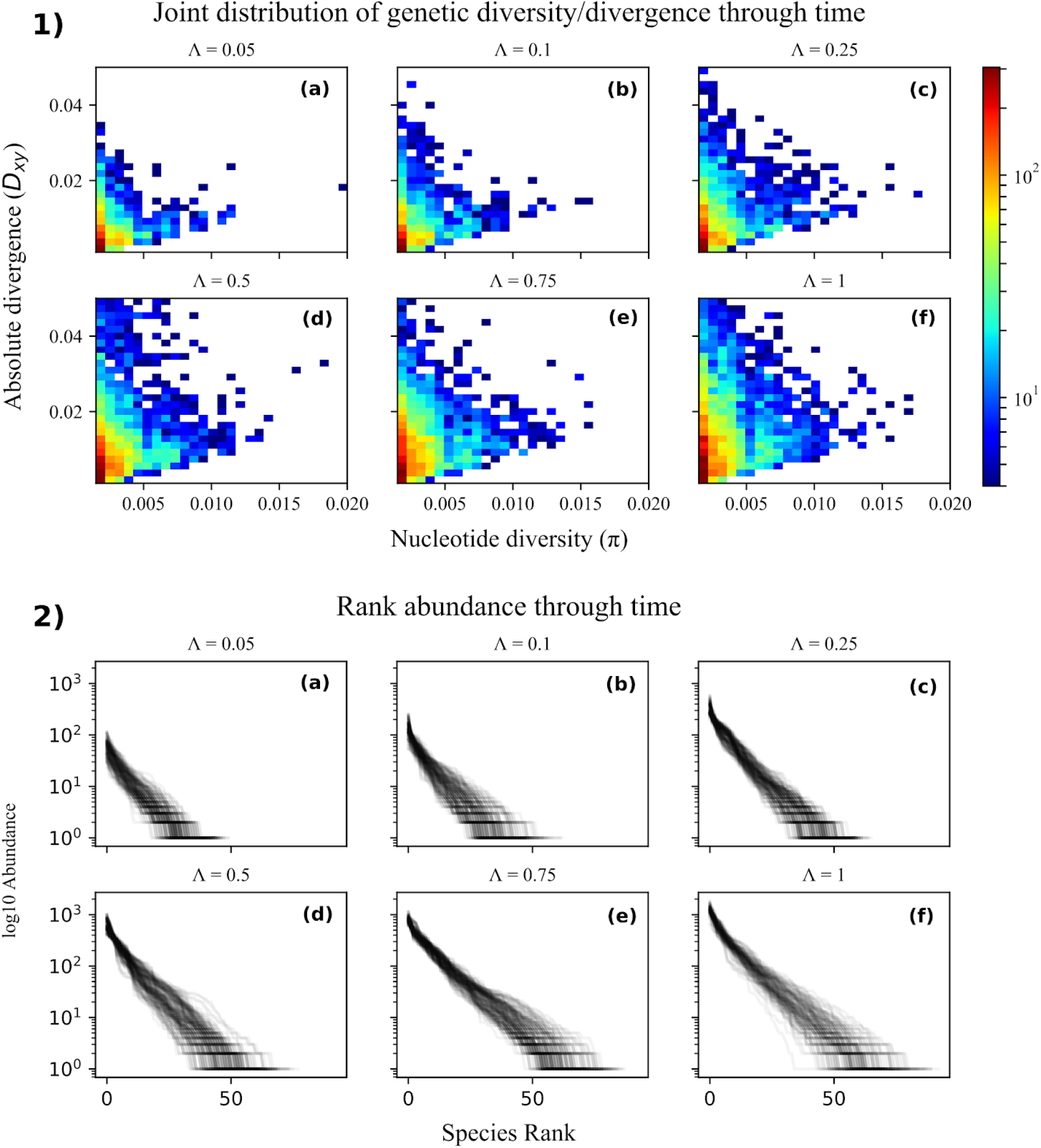
2D-SGD and corresponding rank abundance at varying stages of community assembly. Panel 1) Summed aggregations of the 2D-SGD across 1 × 10^4^ replicated simulations at varying stages of community assembly. All simulations were conducted with intermediate values of community size and colonization rate (K=5000, c=0.03). Each point in the plot is a joint frequency bin for values of local nucleotide diversity (π) and absolute divergence between the local community and the metacommunity (D_xy_). The color of each bin indicates the number of species it contains, with cooler colors signifying fewer species and warmer colors signifying more species. Panel 2) Corresponding rank abundance plots of the 1 × 10^4^ simulated communities. Values of Λ depicted capture multiple stages of community assembly from early (0.05, 0.1), through middle (0.25, 0.5), to late (0.75, 1).

**Figure 3.**
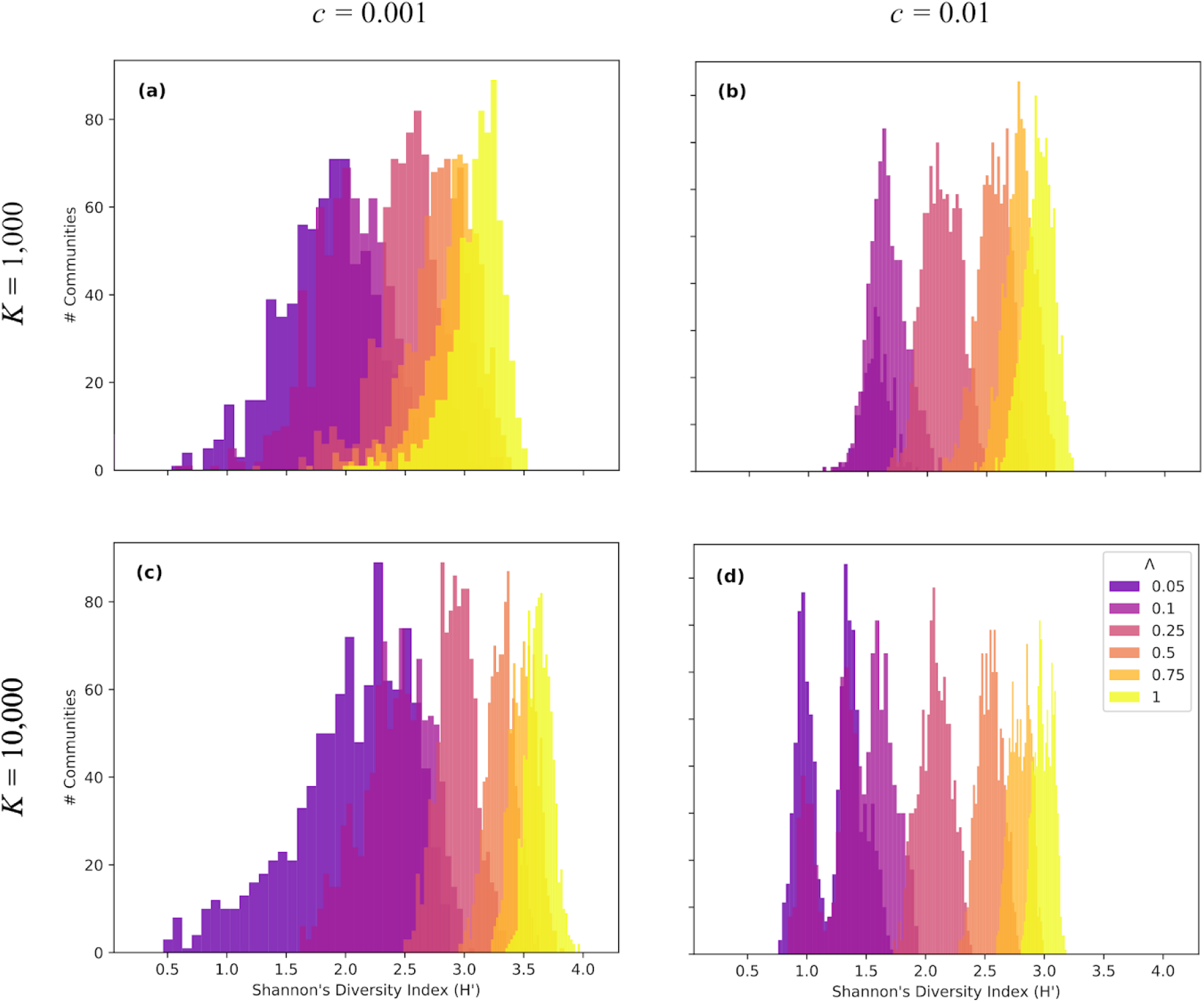
Shannon entropy at varying stages of community assembly. Histograms of Shannon entropy (H’) for four different community parameterizations including low and high colonization, and small and large community sizes. 1 × 10^4^ independent simulations were performed for five Λ values for each parameter combination. Depicted are a) Low colonization rate, small community size; b) High colonization rate, small community size; c) Low colonization rate, large community size; d) High colonization rate, large community size. A range of Λ values were used to capture multiple stages of community assembly from early (0.05, 0.1), through middle (0.25, 0.5), to late (0.75, 1).

Different colonization rates and local community sizes leave different signatures through time on both the SAD and the 2D-SGD. Overall, higher colonization rates tend to increase the species richness in the community, predominantly by increasing the proportion of rare species, as well as species with lower π. Higher colonization also increases local extinction rates (Tables S4 & S5), and this increase in turnover decreases average divergence times, with a subsequent reduction in both n and D_xy_. In a similar fashion, under reduced colonization rates, turnover is lower, the proportion of rare species is reduced, divergence times are longer on average, and n is increased on average (Table S4 & S5). Additionally, the correlation between n and abundance is dependent on A, increasing as A increases, and finally becoming strong as A approaches 1. A powerful feature of our joint model is that it does not assume this correlation between genetic variation and abundance, and indeed the dynamics of how this correlation changes over time provides some of the information for the estimation of model parameters.

### Bias and Accuracy in Estimating Parameters

Broadly speaking, cross-validation indicated reasonable accuracy and limited bias in estimating all parameters under all ABC model configurations (Table S6). The notable exception being estimation under ABC configuration *M_A_*, which is the most data deficient model, as well as when attempting to estimate *K* under all ABC configurations, potentially because the absence of a joint estimate of *K* and A creates identifiability issues. Under ABC model configuration *M*¡, ABC cross-validation indicated a strong signal in the data for estimating H’ using only the 1D-SGD bin values as the summary statistic vector (Fig. 4; RMSE=0.26, R^2^=0.96), with little added value when additionally including D_xy_ under *M_AI_* (Fig. S4; RMSE=0.27, R^2^=0.95). Likewise, A could be estimated well using ABC model configuration *M_I_* and *M_AI_* (R^2^=0.68-72), yet using only H’ as the lone summary statistic (*M_A_*) resulted in poor conditions for estimating Λ (Fig. S4; RMSE=0.28, R^2^=0.05). Our joint framework additionally demonstrated accurate estimation of other ecologically important parameters governing assembly such as community-wide extinction rate (†), and effective colonization rate (c’), with R^2^ between estimated and true values ranging from 0.61-0.89 under ABC model configurations *M_I_*, *M_AI_, M_MI_*, and *M_AMI_* (Table S6; Figs. S4-S7).

**Figure 4.**
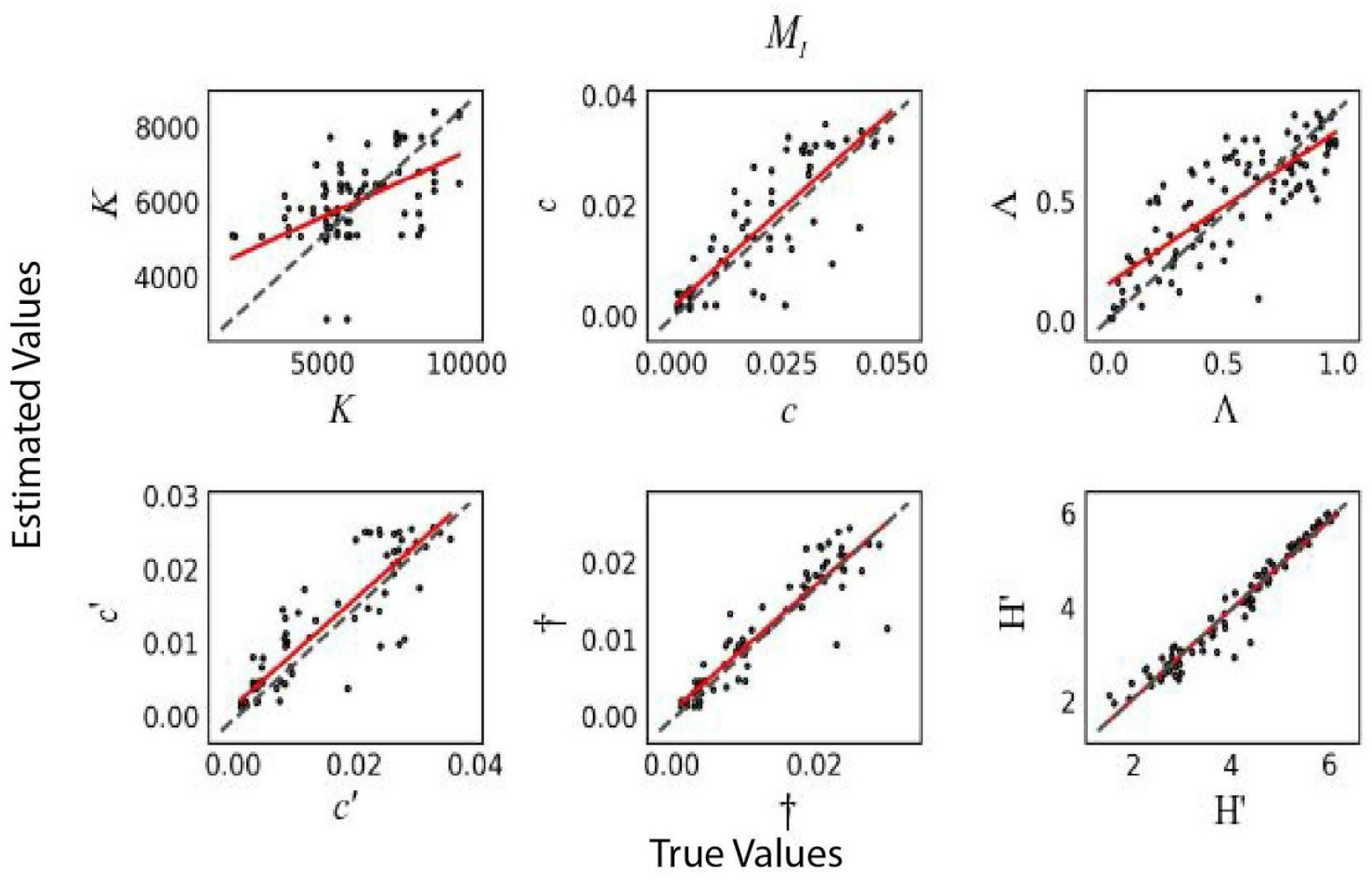
ABC cross-validation for model parameters. 100 ABC cross-validation replicates for comparison of true vs estimated model parameters using only the 1D-SGD as data (*M_I_*). The red line shows the linear least-squares regression between true and estimated values. Results are shown for estimating carrying capacity (*K*), colonization rate (*c*), fraction of equilibrium (Λ), effective colonization rate (*c′*) extinction rate (†), and Shannon entropy (H’).

### Estimating Parameters for the Réunion Spider Community

For an empirical application, we chose to use only the 1D-SGD as observations to estimate H’ calculated from the observed SAD (*M_I_*). In this configuration the bin magnitudes of the 1D-SGD are treated as the summary statistic vector, and H’ is treated as the parameter to be estimated. However, we also have the observed H’ calculated from the samples for direct comparison to the estimate of H’ under the ABC configuration *M_I_*. In this case, our ABC mode estimate of H’ = 1.816 (Fig. 5a; 95% HPD: 1.171-2.822) came remarkably close to the observed H’ of 2.246 calculated from the sampled abundance data. This good fit of the posterior estimate to the observed H’ indicates that the observed distribution of genetic diversity contains sufficient information about the community history of effective population size trajectories across island species with regards to predictions of the contemporary SAD under a neutral model of assembly (Fig. 4). Our simulation study demonstrates this possible dynamic as both H’ and the SGD are predicted to increase over time under most conditions, such that our ABC model could potentially estimate the former with the latter given the strongly temporal features of our assembly model. Given the coupled dynamic of H’ and the SGD as a progressive function of time in our simulation study, it follows that our ABC procedure has potential to estimate the degree of equilibrium parameter (Λ), as shown in our cross-validation experiments. We estimated A for the spider community using three different ABC model configurations configurations representing different combinations of H’ and the 1D-SGD as summary statistics (*M_A_, M_I_*, and *M_AI_*). Given *M_A_* the mode estimate of A was 0.51 but with a diffuse posterior distribution (Fig. 5b; 95% HPD: 0.05-0.93). In sharp contrast, ABC configurations *M_AI_* and *M_I_* yielded mode estimates and HPDs that were both relatively clustered around high values of Λ (Fig. 5c & 5d; posterior mean 0.89; 95% HPD: 0.69-1). Additionally, ABC estimates of *c′* (Fig. 5e; posterior mean 0.001; 95% HPD: 0.0007-0.0017) and f (Fig. 5f; posterior mean 0.001; 95% HPD: 0.0009-0.0012) under model *M_AI_*, were broadly concordant. More formal goodness-of-fit analysis with both the prior predictive check with principal components and Euclidean distances between retained and observed summary statistics corroborate the good fit of the model (Figs. S8 & S9).

**Figure 5.**
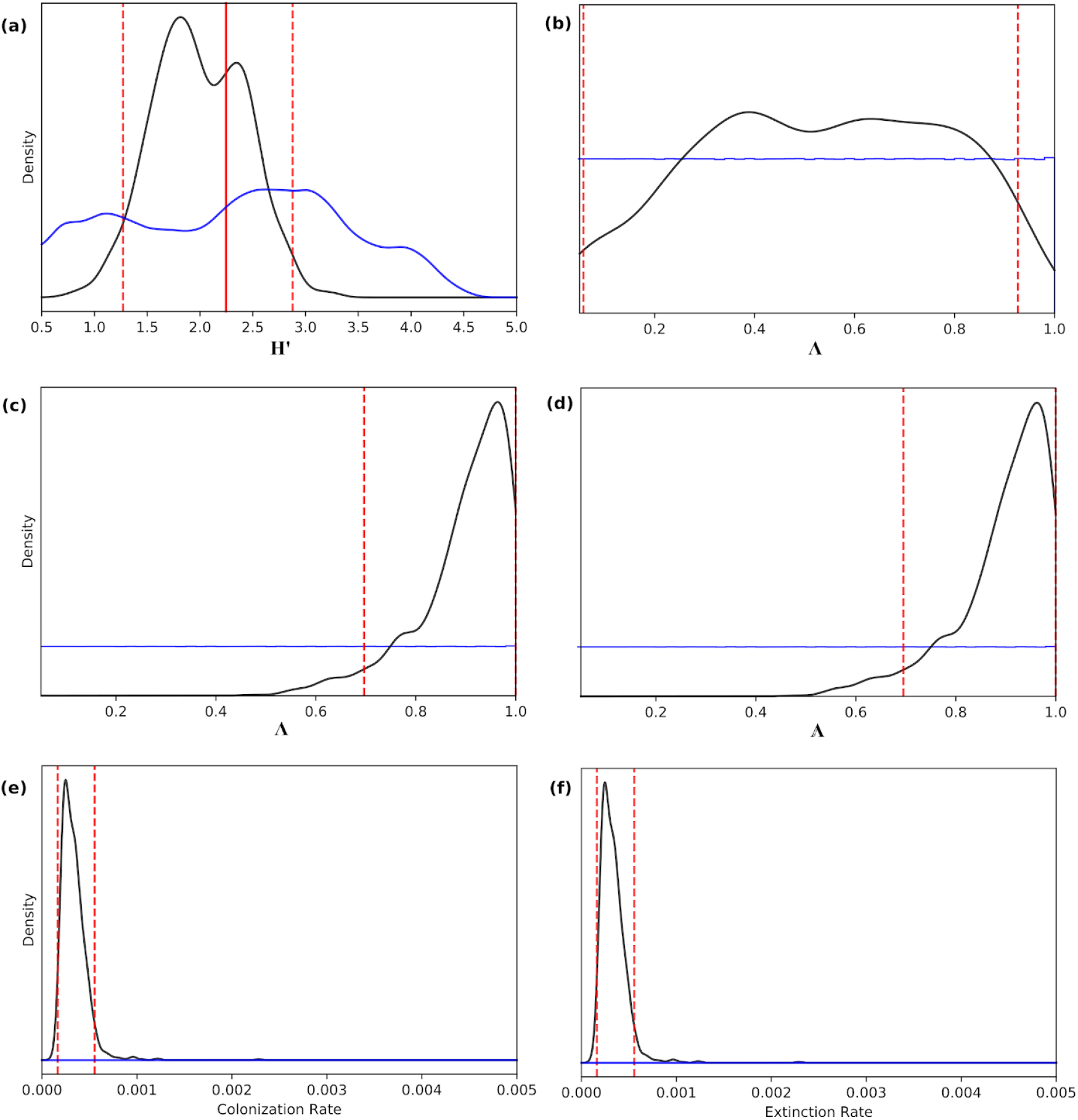
ABC posterior estimates of colonization/extinction rates, H’ and Λ. ABC posterior estimates of colonization and extinction rates, and H’ and Λ for the spider community dataset from the island of Réunion. **(a)** Using only the 1D-SGD as the summary statistic vector (*M_I_*), the mode estimate of H’ was 1.816 (95% HPD: 1.171-2.822; red dashed lines). The true value of H’ as calculated from the observed abundance data was 2.246 (red solid line). Posterior estimates of Λ using three different model configurations: **(b)** only H’ as data (*M_A_*); **(c)** only the 1D-SGD as data (*M_I_*); and **(d)** both H’ and the 1D-SGD as data (*M_AI_*). Posterior estimates of colonization rate and extinction rate using model *M_AI_* are depicted in panels **(e)** and **(f)**, respectively. In all panels the red dashed lines indicate the 95% HPD, and the blue line illustrates the prior distribution.

## Discussion

Similar to efforts that merge phylogenetic frameworks with community assembly models (Jabot & Chave 2009; McPeek 2008; Webb et al. 2002), recent important progress has been made toward linking community ecology models with population genetics (Vellend 2005; Baselga *et al*. 2015; Laroche *et al*. 2015). However, current theory either lacks an explicitly population genetic foundation (Vellend 2005), or considers genetic variation only of a focal taxon (Laroche *et al*. 2015). A focus on genetic diversity at the community scale offers an opportunity for ecological theory to further incorporate the potentially powerful dimension of flexible comparative phylogeographic models (Satler & Carstens 2017; Xue & Hickerson 2017). This should be facilitated by the increasing availability of genome-scale phylogeographic data that allows exploration of evolutionary models of increasing complexity and explanatory power (Schraiber & Akey 2015), yet such approaches have seen limited use to infer the temporal and spatial dynamics at play at the community level (but see Bunnefeld *et al*. 2018). On the other hand, while many classic comparative phylogeographic studies attempted to infer histories of Pleistocene community assembly and diversification (Bermingham & Moritz 1998; Hewitt 2000) by examining combined results of multiple single-taxon phylogeographic studies within a region (Emerson *et al*. 2011), most of these endeavors were not grounded in ecological assembly theory.

Even the comparative phylogeographic models that globally operate at the assemblage level have yet to be grounded in ecological theory that can account for stochastic and deterministic forces underlying community assembly (Prates *et al*. 2016; Gehara *et al*. 2017). Fortunately, the community assembly models that generate expectations for temporally dynamic SADs (Missa *et al*. 2016) and speciation/colonization rates (Rosindell & Harmon 2013) could have an identifiable relationship with population genetic parameters like divergence times, admixture, expansion, colonization times, and changes in effective population sizes. Unifying the parameters of these two modeling frameworks could provide a new way of testing an array of competing assembly models with genetic data as well as estimating the relative strength of various deterministic forces underlying the assembly models such as niche filtering and competition. By linking ecological and micro-evolutionary processes whose dynamics and equilibrium expectations can occur on different time-scales, our new joint approach potentially allows for improved resolution and statistical power for estimating parameters as well as improved potential for testing and fitting a number of different neutral and non-neutral community assembly models. Likewise, understanding whether or not communities tend toward stable equilibria remains an unanswered question (Harmon & Harrison 2015; Rabosky & Hurlbert 2015; Valente *et al*. 2017) that can now be addressed with our joint approach that makes generative predictions of richness, abundance, and the spectrum of genetic diversity under both ecological and evolutionary time scales.

### Assembly of the Réunion Spider Community

The joint data of mitochondrial polymorphism and abundance structure from > 50 spider species on the volcanic island of Réunion affords us the opportunity to compare the estimate of the Shannon entropy (H’) using only the genetic data (i.e. ABC model configuration *M_I_*) with the H’ calculated from the observed abundance distribution. In this case, the posterior distribution of H’ under *M_I_* was able to successfully recover the observed H’. If this is a general feature of our approach it would be encouraging given that estimating species abundances directly from field surveys can be difficult and problematic for some taxa (Kunin *et al*. 2000; Petrovskaya *et al*. 2012).

Using the distributions of abundance and genetic diversity jointly (*M_AI_*) also allowed us to gain insight into the stage of progression towards equilibrium of this spider assemblage under our ecologically neutral model, yet the distribution of genetic diversity alone may have been sufficient (*M_I_*). This was not the case of using the distribution of abundances alone (*M_A_*), indicating there is little information about equilibrium state (Λ) in H’. This result is in agreement with the ABC cross-validation findings suggesting that estimation under ABC configuration *M_AI_* improves accuracy and reduces bias in the estimation of Λ. It is notable that both ABC configurations including local genetic data (*M_I_* and *M_AI_*) strongly indicate that this isolated spider community is consistent with an ecologically neutral assembly that is approaching or has reached equilibrium. Additionally, this assessment is supported by the similar mode estimates and largely overlapping HPD of *c′* and f which hews to the more traditional consideration of equilibrium as the dynamic balance of colonization and extinction. Indeed, Réunion island emerged from a classic volcanic hotspot formation approximately five million years ago (Lénat *et al*. 2001), and this is likely sufficient time for equilibrium expectations of species richness, and community wide distributions of abundance and genetic diversity to have accumulated.

### Outlook

The simple neutral model we introduce can be used as a candidate null hypothesis against which to test comparative population genomic/phylogeographic data, while the flexibility of the framework can accommodate various particular ecological contexts. For example, the model could incorporate *in situ* local speciation either as instantaneous events or as a protracted process (Rosindell *et al*. 2010). Furthermore, it could incorporate non-neutral processes by including trait parameters for differential niche-filtering or dispersal limitation across species that result in variable colonization rates. In this case variation in colonization probabilities would be a proxy for non-neutral processes such as trait-dependent environmental filtering (Pigot & Etienne 2015). Along these lines, the model could also accommodate deterministic processes such as resource-limited colonization probabilities or priority effects while retaining the stochastic dynamics of ecological drift underlying our joint model in the spirit of stochastic assembly theory (Tilman 2004). In this case the magnitude of deviation from neutral expectations of colonization time, abundances, and genetic diversities could be modeled as a free parameter within our joint assembly model.

The increased complexity of these different modelling strategies would all benefit from the increased information content of higher resolution data types such as RADseq (Andrews *et al*. 2016), or whole genomes (Bunnefeld *et al*. 2018). Additionally, the widespread availability of mitochondrial and environmental DNA data also makes our approach amenable to model the assembly of complex microbial systems (Li & Ma 2016) with time-series information (Ridenhour *et al*. 2017). Such time series data could introduce an additional axis of information allowing increased power to test hypotheses about the process of community assembly within a historical perspective.

From a practical standpoint, our model makes it possible to fit assembly models and estimate abundances from a sample of DNA sequence data from a community for which comparable abundance data could be logistically challenging to collect. Taxa with high dispersal potential such as spiders are ideally suited for the estimation of SADs because their genetic samples are more likely to have arisen from a panmictic coalescent process. While taxa with elevated levels of population structure or more complex assembly histories might be more challenging for parameter estimation under our simple model, it could potentially be extended to explicitly model spatial processes, or the more complex assembly histories which may be inherent on real island systems. Our model thus provides a flexible framework that can, even in the absence of comparable species abundance data, allow researchers to use the vast amounts of available mitochondrial DNA sequence data to test competing models of island community assembly.

## Supporting information

## Data accessibility statement

No new data was generated for this study. The joint model implemented in Python and all ipython notebooks for reproducing simulations and analysis are freely available on GitHub: https://github.com/isaacovercast/gimmeSAD.

## Acknowledgements

We wish to thank James Rosindell, Andy Rominger, and three anonymous reviewers for helpful comments on the manuscript. Additionally we thank Silicon Mechanics and their Research Cluster Grant program for the donation of the high-performance computing cluster that was used in support of this research. Funding was provided by grants from FAPESP (BIOTA, 2013/50297-0 to MJH and AC Carnaval), NASA through the Dimensions of Biodiversity Program (DOB 1343578), and the National Science Foundation (DEB-1253710 to MJH). This work would not have been possible without help from the City University of New York High Performance Computing Center, with support from the National Science Foundation (CNS-0855217 and CNS-0958379).

## Author Contributions

IO and MJH conceived the study and designed the model. IO coded the simulations, analyses, and produced the figures. IO wrote the manuscript with the assistance of MJH. BCE contributed empirical data and helped writing the manuscript. All authors reviewed the manuscript.

