## Supplementary material for "An integrated model of population genetics and community ecology"

### Supporting Results

**The Joint SAD and SGD Through Time -** Varying community-wide colonization rate (*c*), community size (*K*), and island origin (volcanic vs continental) also have characteristic impacts on components of the SGD and H' (Fig. S3; Tables S3 & S4). Lower *c* resulted in a greater change in H' over time which was most apparent at lower *K* values and volcanic island settings (Tables S3 & S4), yet H' had the reverse trend under continental island settings. The mean values of $\pi$ and $D_{xy}$ tended to increase over time under the continental island settings whereas only the former tended to increase under the volcanic island setting (Table S3). However, Shannon entropy calculated on these two distributions of genetic diversity ($H'_{\pi}$ .and $H'_{Dxy}$) both tended to increase over time under both island settings. Likewise, the mean values of $\pi$, $D_{xy}$, $H'_{\pi}$ .and $H'_{Dxy}$ tended to all increase over time regardless of the colonization rate (*c*) or community size (*K*), although the magnitude of change depended on these parameter values and island setting (Table S5 & S6).

In the early stages of island community assembly, the 2D-SGDs from volcanic islands differ substantially from those of continental islands (Table S5). There is a priority effect on volcanic islands whereby the initial colonizing species quickly consumes all available ecological space as early arriving populations saturate the local carrying capacity. In this case the initial colonizing species have elevated $\pi$ as well as high $D_{xy}$. However, this genetic signature of the early stage of community assembly quickly erodes as more species gain a foothold on the island, and as $\Lambda$ approaches 1.0 a characteristic distribution of $\pi$ and $D_{xy}$ emerges. In contrast, the early stages of assembly in continental islands are characterized by uniformly higher values of $\pi$ and $D_{xy}$ which tend to decrease as $\Lambda$ approaches 1.0. As $\Lambda$ approaches 1.0, the SAD and 2D-SGD for both island origin models become indistinguishable (Table S5).

**Empirical Analysis Without Samples from the Metacommunity -** Although we do not sample any of the source sister species or sister populations from the mainland, the parameterization of the source meta-community remains under all ABC configurations. A related feature is that our model does not include *in situ* speciation in the local island community, yet because we do not collect data from the source species and do not use any phylogenetic information, *in situ* speciation is perfectly accommodated whereby the formation of new island species from pre-existing island species is parameterized as colonization from the source meta-community.

### Supporting Figures and Tables

#### Table S1. Model Input Parameters

Model parameters, the definition of each parameter, and the values explored in the simulation analyses. The listed parameter values were applied during the ABC analysis of the Réunion spider dataset. For the simulations, the sequence length and infinite sites mutation rate (μ) were chosen to correspond with values for these parameters that are typical for arthropod mitochondrial DNA datasets.

| Parameter | Definition | Values used in simulation experiments |
|---|---|---|
| $K$ | Local community size | ~ uniform(1000-10000) |
| $c$ | Probability an empty deme is replaced by a colonizing individual sampled from the metacommunity (colonization rate) | ~ log-uniform(0.0001-0.01) |
| $\mu$ | Mutation rate | .011 $base^{-1}$ $species^{-1}$ $My^{-1}$ |
| σ | Abundance-coalescence scaling factor | 100 |
| $S_{meta}$ | Number of species in the metacommunity | 1000 |
| $A_{meta}$ | Abundances of species in the metacommunity | ~ logseries(p=0.98) |

**Table S2. Model Response Variables**
Variable names, definitions, and the dimensions of each variable used in the framework.

| Variable | Definition | Dimensions |
|---|---|---|
| $S_{local}$ | Number of species in the local community (i.e. local richness) | Unbounded positive integer |
| $A_i^j$ | Abundance on the island community for species *i* at time *j* | $S_{local} \times \tau^j$ matrix |
| $A_i^j \sigma = N_i^j$ | Time-dependent effective population sizes for species *i* at time *j* | $S_{local} \times \tau^j$ matrix |
| $M_i$ | Post-colonization migrants for species *i* | $S_{local}$ vector |
| $(N_e)_i$ | Harmonic mean of Time-dependent effective population sizes | $S_{local}$ vector |
| $T_i^j$ | Colonization times for species *i* at time *j* | $S_{local}$ vector |
| $A_{meta}$ | Abundances of species in the metacommunity | $S_{meta}$ vector |
| $\Lambda$ | Fraction of equilibrium obtained | Continuous [0, 1] |
| H' | Shannon entropy | Continuous value > 0 |
| † | Effective Extinction rate | Continuous [0, 1] |
| $c'$ | Effective colonization rate | Continuous [0, 1] |

**Table S3. Impact of varying local community origin type**

Mean and standard deviation for Shannon's diversity index of abundances (H'), π, and $D_{xy}$ under either volcanic or continental island origin given K=1000 and c=0.001. These summarizations of abundance and genetic diversity are averaged over 10,000 simulations at five stages of progress toward equilibrium: initial (0.1), early (0.25), moderate (0.5), high (0.75), and complete (1).

| | Mean H' | Stdv H' | Mean $\pi$ | Stdv $\pi$ | Mean $D_{XY}$ | Stdv $D_{XY}$ | Mean $H'_{\pi}$ | Stdv $H'_{\pi}$ | Mean $H'_{Dxy}$ | Stdv $H'_{Dxy}$ |
|---|---|---|---|---|---|---|---|---|---|---|
| | Continental | | | | | | | | | |
| 0.1 | 3.42E+00 | 1.90E-01 | 6.87E-04 | 1.32E-03 | 4.28E-03 | 5.25E-03 | 6.93E-01 | 7.47E-01 | 2.40E+00 | 2.61E-01 |
| 0.25 | 3.27E+00 | 3.15E-01 | 7.53E-04 | 1.90E-03 | 4.51E-03 | 5.84E-03 | 1.10E+00 | 7.56E-01 | 2.73E+00 | 2.49E-01 |
| 0.5 | 2.96E+00 | 2.61E-01 | 9.54E-04 | 1.90E-03 | 4.65E-03 | 5.58E-03 | 1.62E+00 | 6.57E-01 | 3.02E+00 | 1.93E-01 |
| 0.75 | 2.87E+00 | 2.09E-01 | 1.06E-03 | 2.26E-03 | 4.85E-03 | 5.92E-03 | 1.78E+00 | 6.08E-01 | 3.09E+00 | 1.63E-01 |
| 1 | 2.89E+00 | 2.50E-01 | 1.06E-03 | 2.32E-03 | 5.53E-03 | 6.11E-03 | 1.93E+00 | 5.13E-01 | 3.24E+00 | 1.68E-01 |
| | Volcanic | | | | | | | | | |
| 0.1 | 1.17E+00 | 1.26E+00 | 4.31E-04 | 2.95E-04 | 6.28E-03 | 3.05E-03 | 3.37E-01 | 4.40E-01 | 1.16E+00 | 1.15E+00 |
| 0.25 | 2.60E+00 | 6.32E-01 | 3.39E-04 | 1.09E-03 | 5.32E-03 | 5.84E-03 | 7.48E-01 | 4.05E-01 | 2.66E+00 | 5.90E-01 |
| 0.5 | 2.78E+00 | 3.70E-01 | 3.99E-04 | 1.14E-03 | 4.78E-03 | 5.50E-03 | 1.33E+00 | 4.87E-01 | 3.00E+00 | 3.67E-01 |
| 0.75 | 2.75E+00 | 5.17E-01 | 5.45E-04 | 1.61E-03 | 5.48E-03 | 6.01E-03 | 1.43E+00 | 4.37E-01 | 3.05E+00 | 5.23E-01 |
| 1 | 2.87E+00 | 1.23E-01 | 6.30E-04 | 2.01E-03 | 5.40E-03 | 6.22E-03 | 1.67E+00 | 2.49E-01 | 3.21E+00 | 1.55E-01 |

**Table S4. Impact of varying colonization rate**

Mean and standard deviation for Shannon's index (H') and for community level population genetic summary statistics for a volcanic island under varying colonization rates with K=5000. These summarizations of the SAD and of local diversity and island-mainland divergence are averaged over 10,000 simulations at five stages of progress toward equilibrium: initial (0.1), early (0.25), moderate (0.5), high (0.75), and complete (1)

| | Mean H' | Stdv H' | Mean $\pi$ | Stdv $\pi$ | Mean $D_{XY}$ | Stdv $D_{XY}$ | Mean $H'_{\pi}$ | Stdv $H'_{\pi}$ | Mean $H'_{Dxy}$ | Stdv $H'_{Dxy}$ |
|---|---|---|---|---|---|---|---|---|---|---|
| | c = 0.01 | | | | | | | | | |
| 0.1 | 4.25E+00 | 1.15E-01 | 1.53E-04 | 8.67E-04 | 4.44E-03 | 6.22E-03 | 1.60E+00 | 4.02E-01 | 4.06E+00 | 1.55E-01 |
| 0.25 | 4.35E+00 | 9.91E-02 | 1.86E-04 | 9.85E-04 | 4.57E-03 | 6.30E-03 | 2.20E+00 | 3.57E-01 | 4.38E+00 | 1.03E-01 |
| 0.5 | 4.37E+00 | 1.10E-01 | 2.35E-04 | 1.06E-03 | 4.80E-03 | 6.22E-03 | 2.69E+00 | 3.02E-01 | 4.61E+00 | 8.75E-02 |
| 0.75 | 4.42E+00 | 1.28E-01 | 2.78E-04 | 1.11E-03 | 5.08E-03 | 6.29E-03 | 2.98E+00 | 2.61E-01 | 4.73E+00 | 9.28E-02 |
| 1 | 4.39E+00 | 1.06E-01 | 3.41E-04 | 1.32E-03 | 5.60E-03 | 6.84E-03 | 3.12E+00 | 2.36E-01 | 4.79E+00 | 8.43E-02 |
| | c = 0.001 | | | | | | | | | |
| 0.1 | 2.23E+00 | 3.08E-01 | 2.78E-04 | 7.71E-04 | 5.61E-03 | 6.27E-03 | 1.08E+00 | 4.62E-01 | 2.60E+00 | 2.49E-01 |
| 0.25 | 2.32E+00 | 3.28E-01 | 4.67E-04 | 1.17E-03 | 6.83E-03 | 7.34E-03 | 1.60E+00 | 3.84E-01 | 2.84E+00 | 2.06E-01 |
| 0.5 | 2.35E+00 | 2.84E-01 | 7.89E-04 | 1.84E-03 | 8.98E-03 | 9.93E-03 | 1.89E+00 | 3.28E-01 | 2.93E+00 | 1.92E-01 |
| 0.75 | 2.32E+00 | 2.66E-01 | 1.08E-03 | 2.58E-03 | 1.11E-02 | 1.33E-02 | 1.94E+00 | 3.17E-01 | 2.92E+00 | 1.89E-01 |
| 1 | 2.28E+00 | 2.99E-01 | 1.36E-03 | 3.34E-03 | 1.31E-02 | 1.73E-02 | 1.97E+00 | 3.45E-01 | 2.87E+00 | 2.08E-01 |

**Table S5. Impact of varying local community size**

Mean and standard deviation for Shannon's index (H') and for community level population genetic summary statistics for a volcanic island under varying sizes with fixed c=0.001. These summarizations of the SAD and of local diversity and island-mainland divergence are averaged over 10,000 simulations at five stages of progress toward equilibrium: initial (0.1), early (0.25), moderate (0.5), high (0.75), and complete (1)

| | Mean H' | Stdv H' | Mean $\pi$ | Stdv $\pi$ | Mean $D_{XY}$ | Stdv $D_{XY}$ | Mean $H'_{\pi}$ | Stdv $H'_{\pi}$ | Mean $H'_{Dxy}$ | Stdv $H'_{Dxy}$ |
|---|---|---|---|---|---|---|---|---|---|---|
| | K = 1000 | | | | | | | | | |
| 0.1 | 9.98E-01 | 4.28E-01 | 3.00E-04 | 4.73E-04 | 6.58E-03 | 5.72E-03 | 1.74E-01 | 3.03E-01 | 1.17E+00 | 3.88E-01 |
| 0.25 | 9.98E-01 | 4.23E-01 | 5.93E-04 | 9.48E-04 | 7.84E-03 | 6.84E-03 | 3.77E-01 | 4.01E-01 | 1.32E+00 | 3.88E-01 |
| 0.5 | 9.30E-01 | 4.77E-01 | 9.41E-04 | 1.50E-03 | 9.24E-03 | 8.07E-03 | 5.13E-01 | 4.33E-01 | 1.41E+00 | 4.08E-01 |
| 0.75 | 9.52E-01 | 4.56E-01 | 1.49E-03 | 2.25E-03 | 1.11E-02 | 1.02E-02 | 6.47E-01 | 4.27E-01 | 1.39E+00 | 3.96E-01 |
| 1 | 1.09E+00 | 3.48E-01 | 1.89E-03 | 2.55E-03 | 1.08E-02 | 1.04E-02 | 8.85E-01 | 2.86E-01 | 1.53E+00 | 3.27E-01 |
| | K = 5000 | | | | | | | | | |
| 0.1 | 2.23E+00 | 3.08E-01 | 2.78E-04 | 7.71E-04 | 5.61E-03 | 6.27E-03 | 1.08E+00 | 4.62E-01 | 2.60E+00 | 2.49E-01 |
| 0.25 | 2.32E+00 | 3.28E-01 | 4.67E-04 | 1.17E-03 | 6.83E-03 | 7.34E-03 | 1.60E+00 | 3.84E-01 | 2.84E+00 | 2.06E-01 |
| 0.5 | 2.35E+00 | 2.84E-01 | 7.89E-04 | 1.84E-03 | 8.98E-03 | 9.93E-03 | 1.89E+00 | 3.28E-01 | 2.93E+00 | 1.92E-01 |
| 0.75 | 2.32E+00 | 2.66E-01 | 1.08E-03 | 2.58E-03 | 1.11E-02 | 1.33E-02 | 1.94E+00 | 3.17E-01 | 2.92E+00 | 1.89E-01 |
| 1 | 2.28E+00 | 2.99E-01 | 1.36E-03 | 3.34E-03 | 1.31E-02 | 1.73E-02 | 1.97E+00 | 3.45E-01 | 2.87E+00 | 2.08E-01 |

**Table S6. ABC cross-validation results**

Root mean squared error (RMSE) and coefficient of determination ($R^2$) for 100 ABC leave-one-out"cross-validation replicates for each model parameter under each of the five data availability scenarios. Parameters investigated include carrying capacity (K), parameterized colonization rate (c), fraction of equilibrium (Λ), calculated colonization rate (c'), extinction rate (†), and Shannon's diversity index (H').

| | $M_A$ | | $M_I$ | | $M_{AI}$ | | $M_{MI}$ | | $M_{AMI}$ | |
|---|---|---|---|---|---|---|---|---|---|---|
| | RMSE | $R^2$ | RMSE | $R^2$ | RMSE | $R^2$ | RMSE | $R^2$ | RMSE | $R^2$ |
| K | 1761 | 0.15 | 1409 | 0.33 | 1371 | 0.43 | 1774 | 0.45 | 1595 | 0.6 |
| c | 0.07 | 0.83 | 0.007 | 0.78 | 0.005 | 0.87 | 0.006 | 0.82 | 0.004 | 0.87 |
| Λ | 0.28 | 0.05 | 0.18 | 0.68 | 0.17 | 0.72 | 0.18 | 0.67 | 0.18 | 0.6 |
| c' | 0.004 | 0.8 | 0.005 | 0.82 | 0.006 | 0.71 | 0.005 | 0.78 | 0.004 | 0.84 |
| † | 0.003 | 0.89 | 0.003 | 0.89 | 0.004 | 0.76 | 0.003 | 0.83 | 0.003 | 0.89 |
| H' | | | 0.26 | 0.96 | | | 0.27 | 0.95 | | |

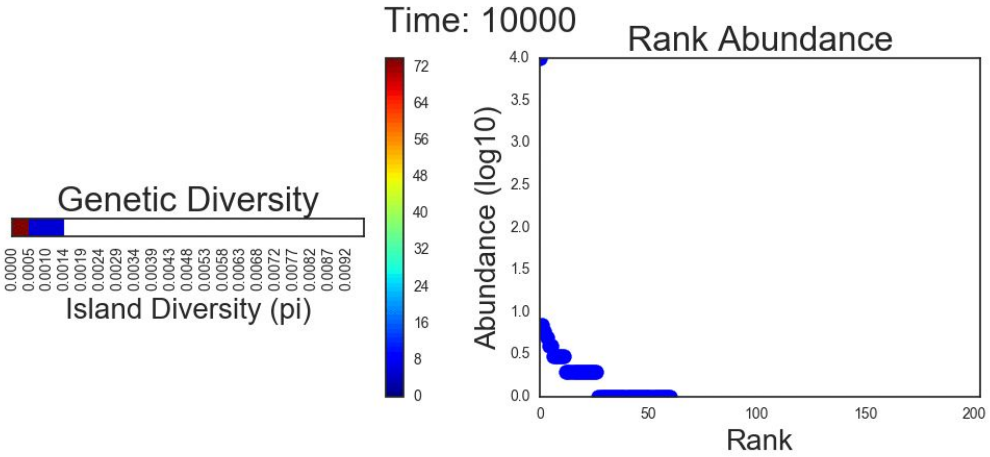


**Figure S1. One time step of a simulated 1D-SGD**

One time step of a simulated 1D-SGD plotted beside the associated rank abundance distribution of the local community from which it was generated. Left panel: 1-dimensional histogram illustrating the community level genetic profile (forward in time). Each point in the plot is a frequency bin for values of local nucleotide diversity ($\pi$). The color of each bin indicates the number of species it contains, with cooler colors signifying fewer species and warmer colors signifying more species. Right panel: Rank abundance plot of the simulated community at the same time-point. Points in the plot are species ranked by their abundance in the local community. Note that the Y-axis (abundance) is log scaled. Time is indicated in generations of the forward time assembly model. A complete realization of the 1D-SGD is included as a video in the electronic supporting information.

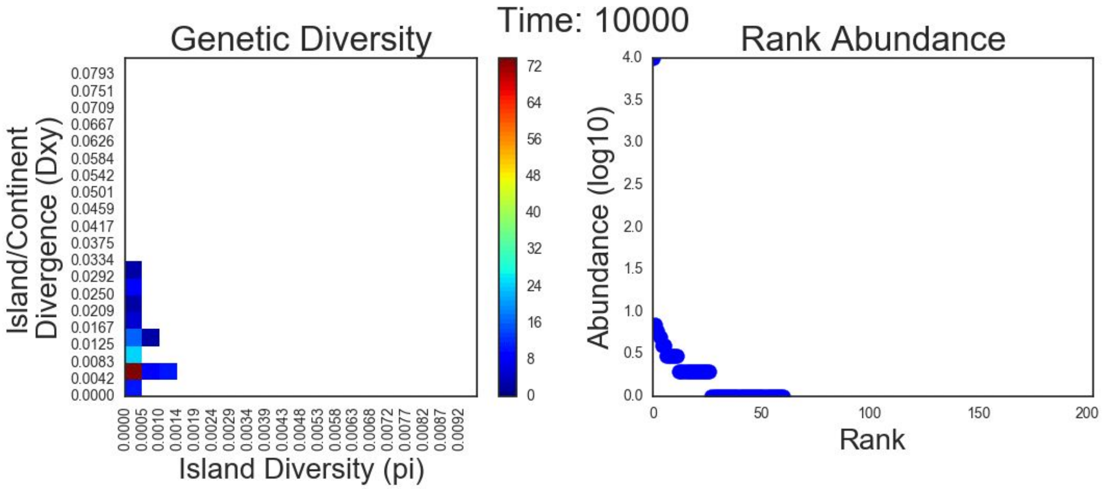


**Figure S2 - One time step of a simulated 2D-SGD**

One time step of a simulated 2D-SGD plotted beside the associated rank abundance distribution of the local community from which it was generated. Left panel: 2-dimensional histogram illustrating the community level genetic profile (forward in time). Right panel: Rank abundance plot of the simulated community at the same time-point. Note that the Y-axis (abundance) is log scaled. Time is indicated in generations of the forward time assembly model. A complete realization of the 2D-SGD is included as a video in the electronic supporting information.

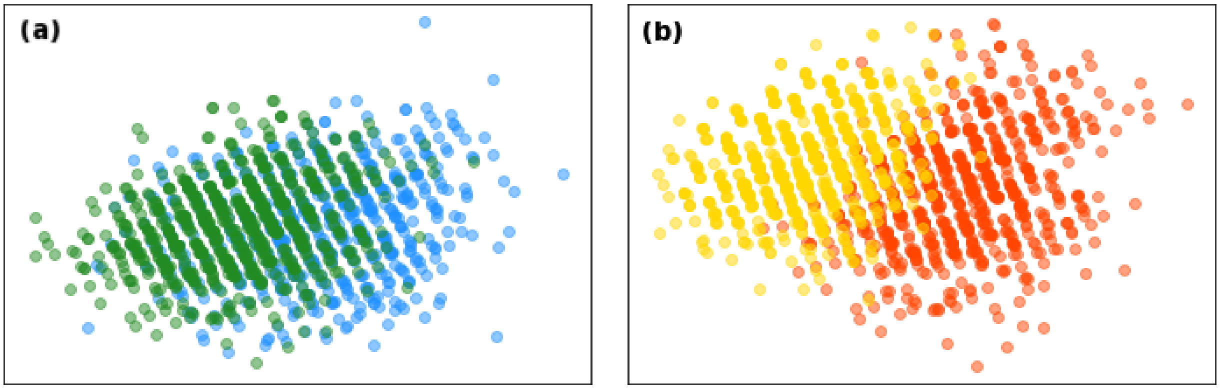


**Figure S3. PCA of 1000 simulated 2D-SGD for different model parameterizations**
PCA plots of $1x10^3$ simulated joint frequencies of community level $\pi$ and $D_{xy}$ (2D-SGD) at $\Lambda = 0.5$ for 4 different simulation conditions. **(a)** Two models under K=1000 with $c$ = 0.001 (blue) and $c$ = 0.01 (green) colonization rates. **(b)** Two models under K=5000 and $c$ = 0.001 (yellow) and $c$ = 0.01 (red) colonization rates. Both plots depict PC1 on the X-axis and PC2 on the Y-Axis.

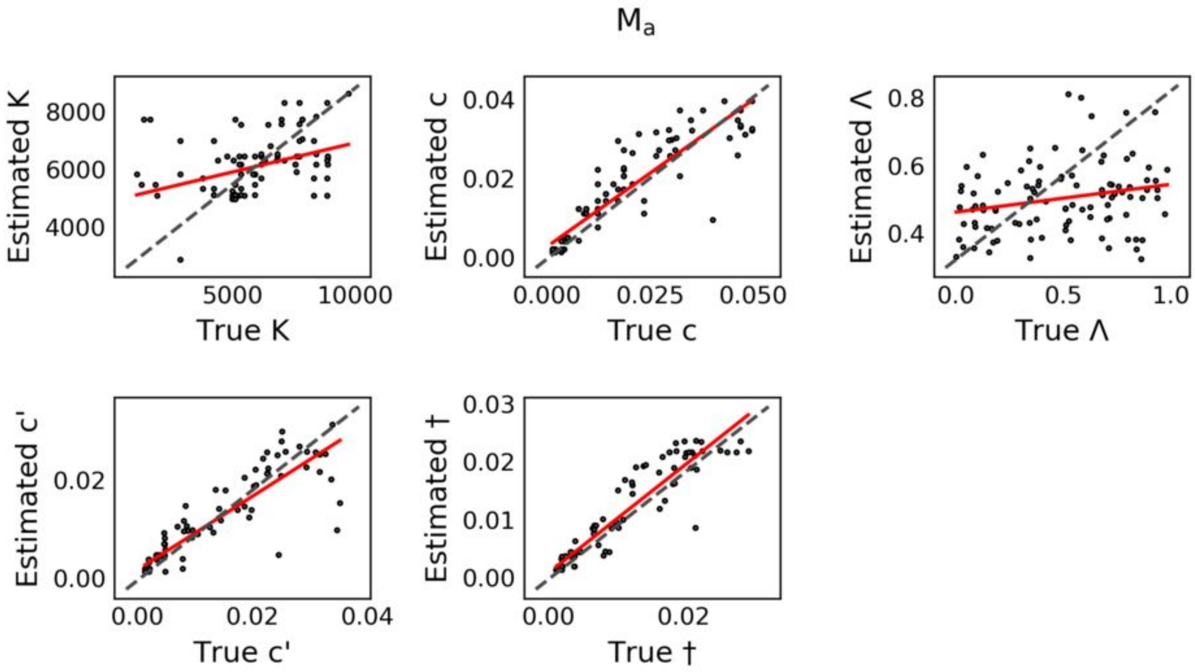


**Figure S4. ABC cross-validation for parameter estimates with ABC configuration $M_A$**
100 ABC cross-validation replicates for estimating model parameters using only abundances as data ( $M_A$ ). The identity line is plotted with a dashed line. The red line shows the linear least-squares regression between true and estimated values. Results are shown for carrying capacity (K), parameterized colonization rate (c), fraction of equilibrium (Λ), calculated colonization rate (c'), extinction rate (†), and Shannon's diversity index (H').

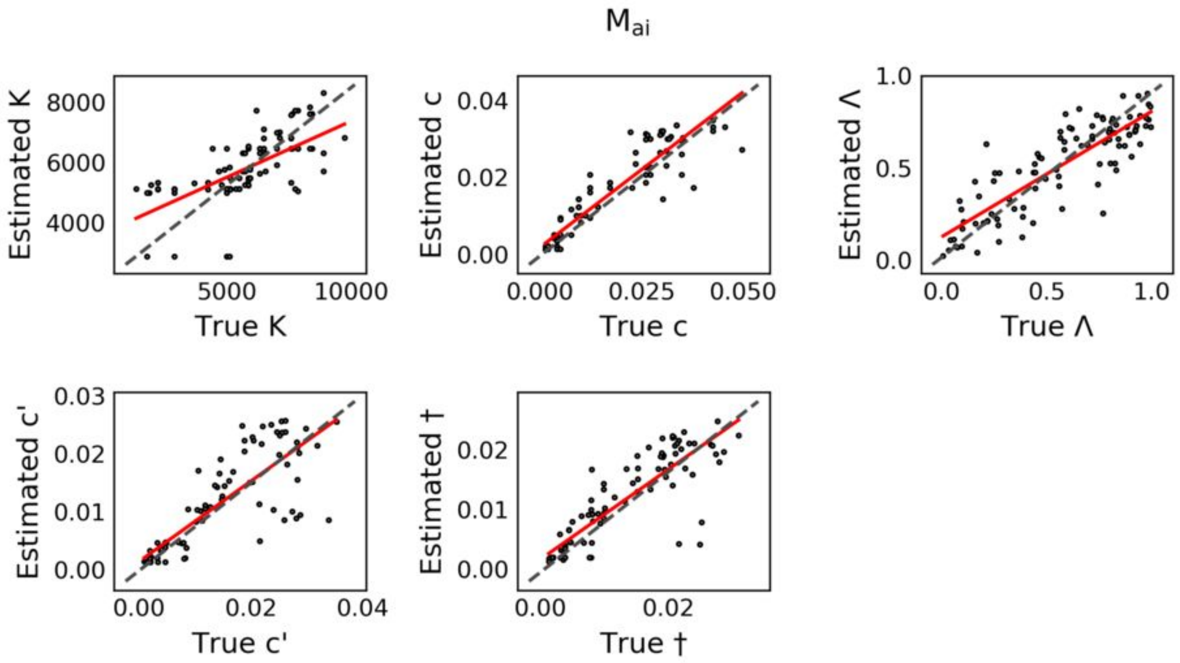


**Figure S5. ABC cross-validation for parameters estimates with ABC configuration** $\boldsymbol{M_{AI}}$
100 ABC cross-validation replicates for estimating model parameters using both abundances and the 1D-SGD as data ( $M_{AI}$ ). The identity line is plotted with a dashed line. The red line shows the linear least-squares regression between true and estimated values. Results are shown for carrying capacity (K), parameterized colonization rate (c), fraction of equilibrium (Λ), calculated colonization rate (c'), extinction rate (†), and Shannon's diversity index (H').

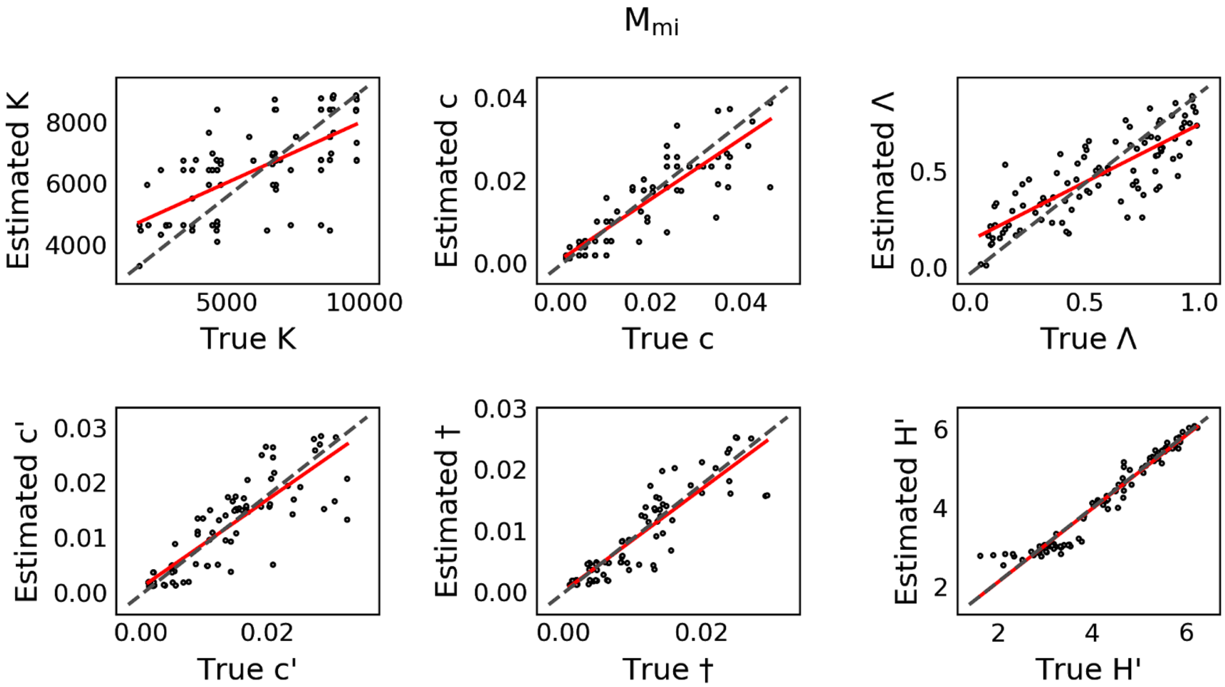


**Figure S6. ABC cross-validation for parameters estimates with ABC configuration $M_{MI}$**
100 ABC cross-validation replicates for estimating model parameters using the 2D-SGD as data ($M_{MI}$). The identity line is plotted with a dashed line. The red line shows the linear least-squares regression between true and estimated values. Results are shown for carrying capacity (K), parameterized colonization rate (c), fraction of equilibrium (Λ), calculated colonization rate (c'), extinction rate (†), and Shannon's diversity index (H').

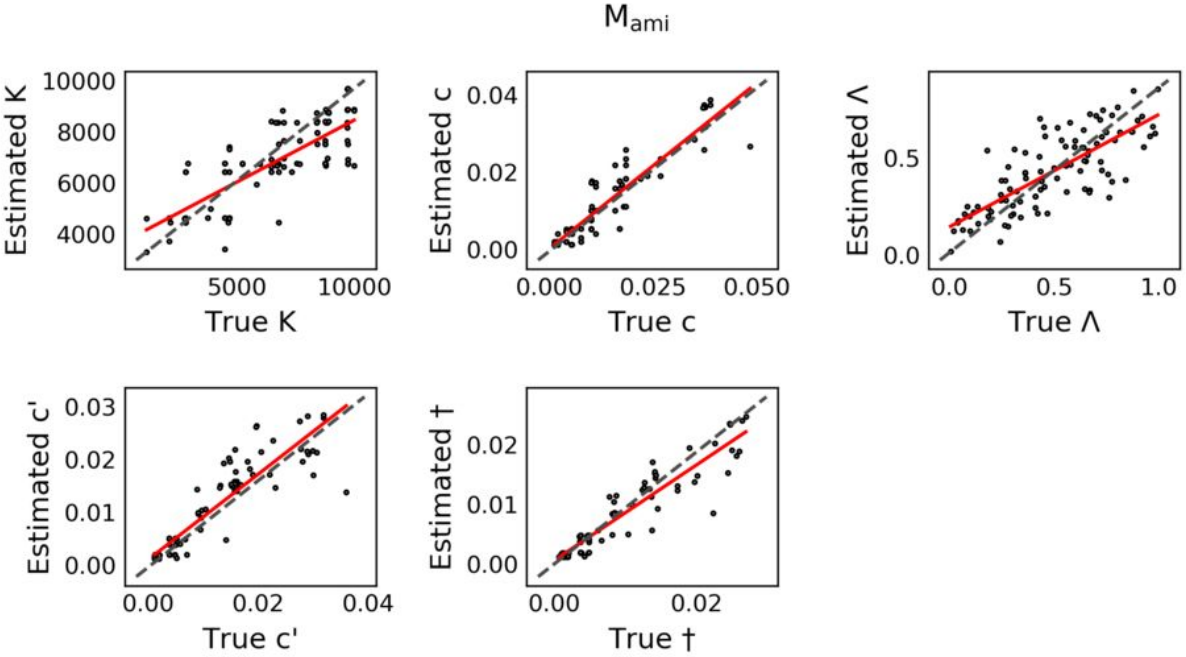


**Figure S7. ABC cross-validation for parameters estimates with ABC configuration $M_{AMI}$**
100 ABC cross-validation replicates for estimating model parameters using both abundances and the 2D-SGD as data ($M_{AMI}$). The identity line is plotted with a dashed line. The red line shows the linear least-squares regression between true and estimated values. Results are shown for carrying capacity (K), parameterized colonization rate (c), fraction of equilibrium (Λ), calculated colonization rate (c'), extinction rate (†), and Shannon's diversity index (H').

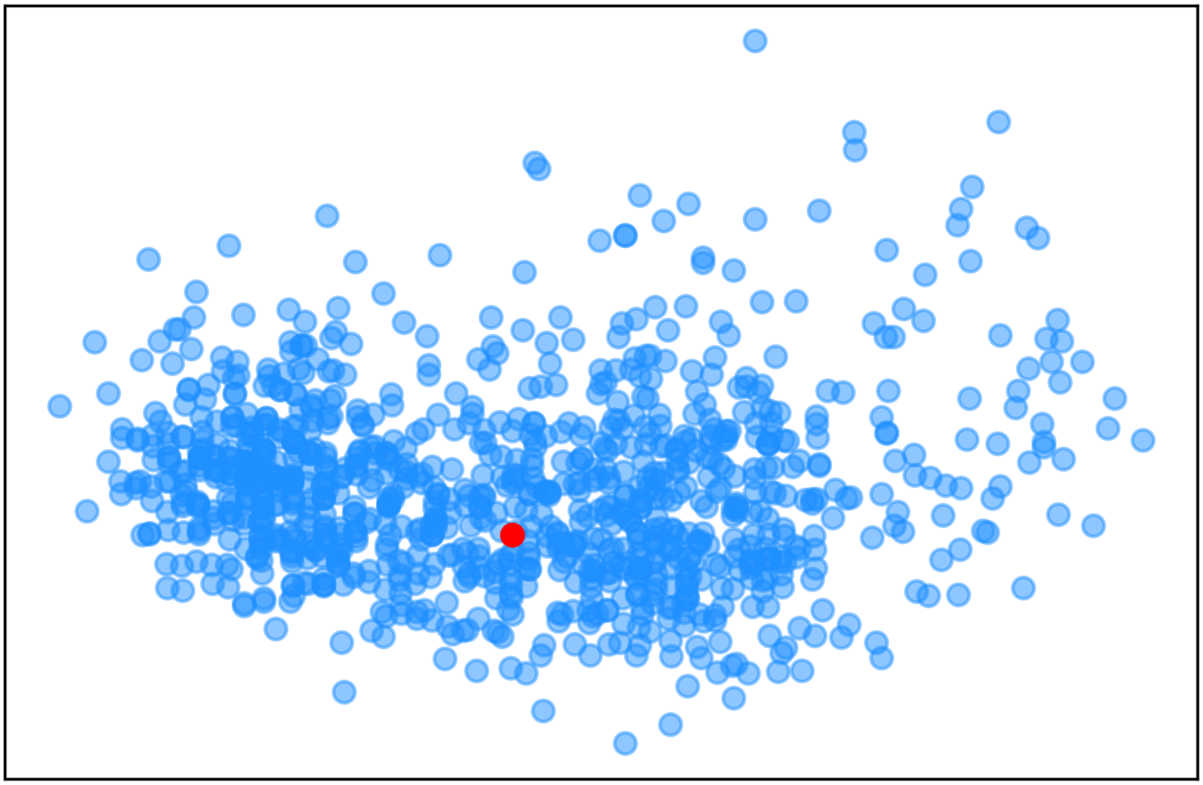

**Figure S8. PCA of retained simulations with empirical data**
Prior predictive check for goodness of fit of retained simulations to the empirical data. All 1000 retained simulated 1D-SGD are shown projected into the first two axes of principal component space, along with the 1D-SGD calculated from the observed Réunion spider data (in red).

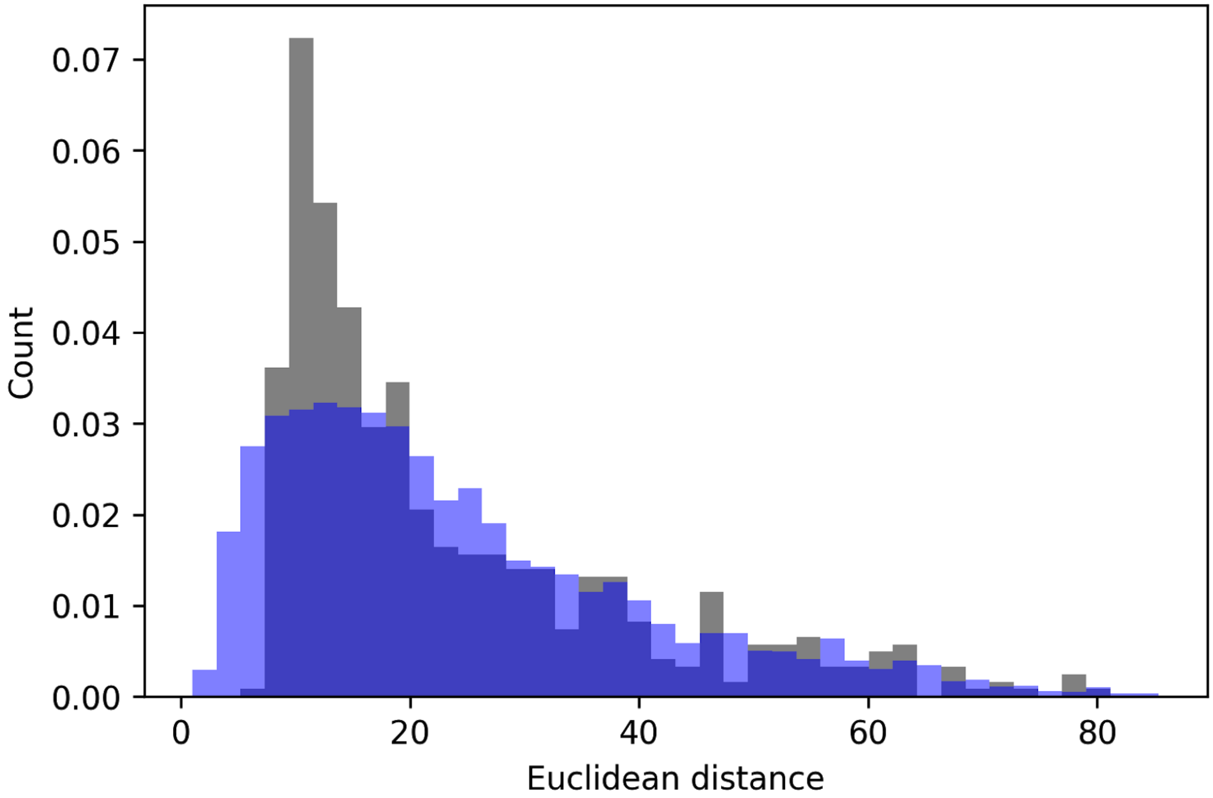


**Figure S9. Euclidean distance between observed and estimated diversity**
Euclidean distance between the 1D-SGD of the 1000 retained simulations from ABC analysis and the 1D-SGD calculated from the observed data (grey). Also shown is the distribution of pairwise Euclidean distances among all 1D-SGD of the retained simulations (blue). Distances among retained simulations are plotted with reduced alpha so that characteristics of both distributions can be observed simultaneously.

**Box S1:** gimmeSAD model overview

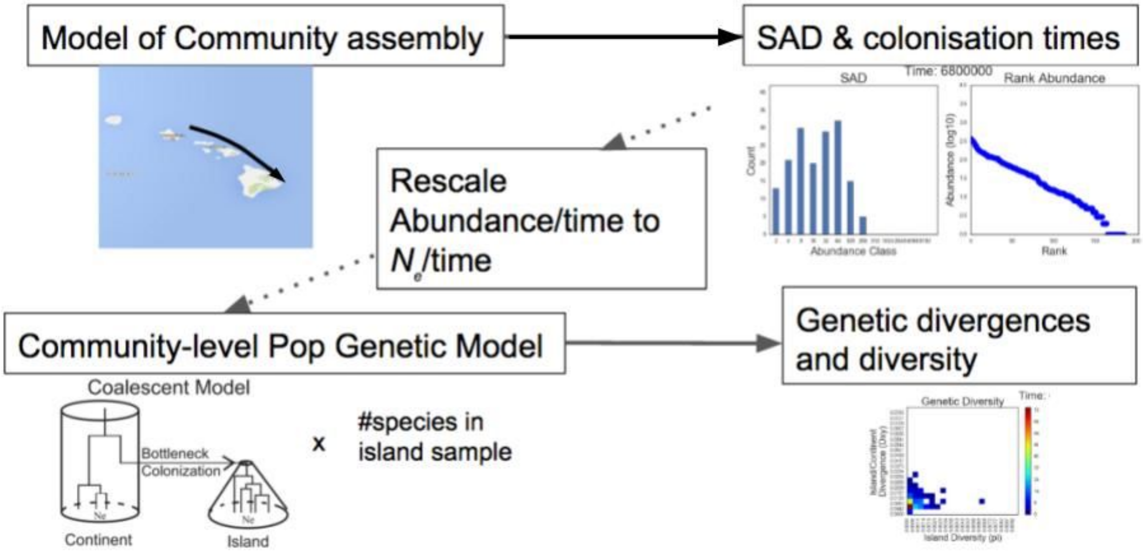


1) Forward time simulations of community assembly follow the spatially implicit neutral model of whereby abundance distributions, and immigration and extinction rates proceed under a birth/death/colonization process. The carrying capacity (K) of the local community consists of the sum of population sizes of all species on the island, and the colonization rate is modeled as a single parameter (c) that specifies the probability of a colonization event from the metacommunity into the local community.
2) At each time-step one individual is randomly sampled for removal from the local community. With probability 1-c this individual is replaced by the offspring of a randomly sampled individual from the local community. With probability c, the individual is replaced by a randomly sampled member of the metacommunity.
3) Local abundances and colonization times are tracked for all species extant in the local community for each forward time epoch, from initial colonization up to and then through equilibrium.
4) For each forward time epoch, for each extant species, local abundance and colonization time are rescaled to effective population size ($N_e$) and divergence time. Sequence data is simulated using a backward time coalescent model parameterized by these values, and local diversity ($\pi$) and island/metacommunity divergence ($D_{xy}$) are calculated.
5) Finally, $\pi$ and $D_{xy}$ values are binned to generate the joint frequency histogram which summarizes community level patterns of genetic diversity and divergence.
